# Long-Term Inactivation of Sodium Channels as a Mechanism of Adaptation in CA1 Pyramidal Neurons

**DOI:** 10.1101/2021.10.26.465936

**Authors:** Carol Upchurch, Crescent Combe, Christopher Knowlton, Valery Rousseau, Sonia Gasparini, Carmen C. Canavier

**Author notes:** Corresponding author: Sonia Gasparini, Neuroscience Center, Louisiana State University Health Sciences Center, New Orleans, 2020 Gravier St., New Orleans, LA 70112. these authors contributed equally (Carol Upchurch, Crescent Combe). these authors jointly supervised this work (Sonia Gasparini, Carmen C. Canavier).

## Abstract

Many hippocampal CA1 pyramidal cells function as place cells, increasing their firing rate when a specific place field is traversed. The dependence of CA1 place cell firing on position within the place field is asymmetric. We investigated the source of this asymmetry by injecting triangular depolarizing current ramps to approximate the spatially-tuned, temporally-diffuse depolarizing synaptic input received by these neurons while traversing a place field. Ramps were applied to CA1 pyramidal neurons from male rats *in vitro* (slice electrophysiology) and *in silico* (multi-compartmental NEURON model). Under control conditions, CA1 neurons fired more action potentials at higher frequencies on the up-ramp versus the down-ramp. This effect was more pronounced for dendritic compared to somatic ramps. We incorporated a four-state Markov scheme for Na_V_1.6 channels into our model and calibrated the spatial dependence of long-term inactivation according to the literature; this spatial dependence was sufficient to explain the difference in dendritic versus somatic ramps. Long-term inactivation reduced the firing frequency by decreasing open-state occupancy, and reduced spike amplitude during trains by decreasing occupancy in closed states, which comprise the available pool. PKC activator phorbol-dibutyrate, known to reduce Na_V_ long-term inactivation, removed spike amplitude attenuation *in vitro* more visibly in dendrites and greatly reduced adaptation, consistent with our hypothesized mechanism. Intracellular application of a peptide inducing long-term Na_V_ inactivation elicited spike amplitude attenuation during spike trains in the soma and greatly enhanced adaptation. Our synergistic experimental/computational approach shows that long-term inactivation of Na_V_1.6 is a key mechanism of adaptation in CA1 pyramidal cells.

**Significance statement:** The hippocampus plays an important role in certain types of memory, in part through context-specific firing of “place cells”; these cells were first identified in rodents as being particularly active when an animal is in a specific location in an environment, called the place field of that neuron. In this in vitro/in silico study, we found that long-term inactivation of sodium channels causes adaptation in the firing rate that could potentially skew the firing of CA1 hippocampal pyramidal neurons earlier within a place field. A computational model of the sodium channel revealed differential regulation of spike frequency and amplitude by long-term inactivation, which may be a general mechanism for spike frequency adaptation in the central nervous system.

## Introduction

Area CA1 of the hippocampus is thought to play a key role in learning and memory, specifically episodic memories (Burgess et al., 2002), sequential order (Fortin et al., 2002; Hoang and Kesner, 2008) and trace conditioning (Shors, 2004; Hunsaker et al., 2006). A type of episodic memory involving the sequential order of places along a linear track has been well studied in rodents (O’Keefe and Nadel, 1978). Many CA1 pyramidal cells function as place cells (O’Keefe and Dostrovsky, 1971); they increase their firing rate as they traverse a place field. These fields are specific to a given environment (Colgin et al., 2008), and result from the activation of a particular subset of synaptic inputs by aspects of the place field in that environment. The firing rate of individual place cells is generally symmetric around a peak in the center of the place field, as recorded in head-restrained mice navigating a virtual spatial environment on a spherical treadmill (Harvey et al., 2009). However, intracellular recordings from the same mice revealed that traversing the place field evoked a depolarizing ramp of synaptic excitation that was not symmetric, but instead peaked after three quarters of the field had been traversed, such that the peaks of the firing rate and the underlying depolarization do not coincide. One explanation for this asymmetry is provided by the response of CA1 neurons to a temporally symmetric current ramp in anesthetized mice (Harris et al., 2002). The firing rate adapts such that the neuron fires substantially less on the down-ramp compared to the up-ramp. Such adaptation can explain how the firing rate peaks before the synaptic depolarizing ramp peaks. Although Harvey et al. (2009) observed a symmetric dependence of the firing rate on distance through the place field, this dependence becomes skewed in a predictive direction as the center of mass moves earlier in the place field as the environment becomes more familiar (Mehta et al., 2000), likely through synaptic plasticity mechanisms. Since adaptation is likely to contribute to the skew of the firing rate, here we investigate intrinsic mechanisms that determine firing rate adaptation in CA1 pyramidal cells.

Adaptation can result from accumulation of an outward current or a decrement in an inward current active in the inter-spike interval. The Ca^2+^-activated small-conductance K^+^ current (I_SK_) (Stocker et al., 1999; Pedarzani et al., 2005) and the muscarinic M-type K^+^ current (I_M_) (Otto et al., 2006; Gu et al., 2008) are often implicated in firing rate adaptation in CA1 pyramidal neurons. Voltage-gated Na^+^ channels in CA1 pyramidal neurons have previously been shown to exhibit long-term inactivation that increases with distance from the soma along the apical dendrite (Colbert et al., 1997; Jung et al., 1997), and the resulting decrease in the available pool of sodium channels has been suggested as a mechanism for frequency adaptation in the same neurons (Fernandez and White, 2010).

We systematically tested these possible mechanisms of adaptation using a combination of *in vitro* slice electrophysiology and computational modeling. We applied symmetric current ramps to the soma or apical dendrite of CA1 pyramidal cells to simulate the depolarizing synaptic input these cells receive *in vivo* while crossing a place field. Selective blockers of I_SK_ and I_M_ did not decrease adaptation. To investigate the putative contribution of long-term inactivation of Na^+^ channels to adaptation, we calibrated a Markov model of a Na^+^ channel with a second inactivated state that recovered more slowly than the first, according to the spatially-dependent long-term inactivation shown in Mickus et al. (1999).

Experimentally, we observed that adaptation was more pronounced in the dendrites compared to the soma. The model demonstrated that the increase in occupancy in the long-term inactivated state of sodium channels along the apical dendrite was sufficient to account for this difference. Increasing or decreasing the occupancy in the long-term inactivated state, through a selective peptide or PKC activator, respectively, significantly enhanced or attenuated adaptation, as predicted by the model.

## Materials and Methods

### Experimental methods

#### Slice preparation

All the procedures described were conducted according to protocols approved by the Louisiana State University Health Sciences Center-New Orleans Institutional Animal Care and Use Committee, following guidelines on the responsible use of laboratory animals in research from the National Institutes of Health. 7 to 11-week-old male Sprague Dawley rats were deeply anesthetized via intraperitoneal injection of ketamine and xylazine (90 and 10 mg/kg, respectively). Once the toe-pinch and palpebral reflexes ceased, rats were transcardially perfused with ice-cold oxygenated cutting solution containing, in mM: NaHCO_3_ 28, KCl 2.5, NaH_2_PO_4_ 1.25, MgCl_2_ 7, CaCl_2_ 0.5, dextrose 7, sucrose 234, L-ascorbic acid 1, sodium pyruvate 3, and decapitated. The brains were rapidly removed and a vibratome used to cut 400 µm-thick transverse hippocampal slices that were then transferred to a chamber filled with an oxygenated artificial cerebro-spinal fluid (ACSF) containing, in mM: NaCl 125, NaHCO_3_ 25, KCl 2.5, NaH_2_PO_4_ 1.25, MgCl_2_ 1, CaCl_2_ 2, dextrose 25, ascorbate 1, sodium pyruvate 3. After the cutting procedure, slices were allowed to recover for one hour at 36°C, with an additional recovery period of at least one hour at room temperature.

#### Patch clamp electrophysiology

Individual slices were transferred to a submerged recording chamber, and superfused with ACSF. All experiments were performed at 34-36°C. CA1 pyramidal cells were identified via differential interference contrast-infrared video microscopy. Whole-cell current clamp recordings were made using Dagan BVC 700A amplifiers in the active “bridge” mode. Recording pipettes had a resistance of 1-3 MΩ (for somatic recordings) and 3-5 MΩ (for dendritic recordings) when filled with an internal solution that contained, in mM: potassium methanesulphonate 125, KCl 20, HEPES 10, EGTA 0.5, NaCl 4, Mg_2_ATP 4, Tris_2_GTP 0.3, phosphocreatine 14. Cells with resting membrane potentials depolarized beyond −60 mV at break-in were discarded. Series resistance was monitored throughout the recordings and was usually less than 20 MΩ; recordings were discarded when, series resistance reached 25 MΩ or 30 MΩ, for somatic and dendritic recordings, respectively.

NBQX (10 µM), DL-APV (50 µM), Gabazine (12.5 µM) and CGP55845 (1 µM) were applied in the external solution in all experiments to block glutamatergic and GABAergic neurotransmission, respectively, in order to isolate the contribution of the intrinsic ion channels to the ramp responses. Apamin (100 nM), XE-991 (10 µM), phorbol-2,13-dibutyrate (PDBu, 5 µM), and *Androctonus mauretanicus mauretanicus toxin 3 (*AmmTx3, 300 nM) were added to ACSF as needed, from stock solutions made with water or DMSO; the concentration of DMSO in the final solution was ≤ 0.1%. In a set of experiments to induce long-term inactivation of Na_V_ channels, an N-terminally acetylated peptide, corresponding to residues 2-21 of the fibroblast growth factor homologous factor 2A (FHF2A, Dover et al., 2010) was added to the pipette intracellular solution. The peptide (F2A(2-21)) was purchased from Thermo Fisher Scientific (Rockford, IL) where it was custom synthesized, purified by HPLC and its identity confirmed by mass spectroscopy. Gabazine and CGP55845 were purchased from Abcam (Cambridge, MA), DL-APV and XE-991 were from HelloBio (Princeton, NJ), NBQX, apamin and AmmTx3 were from Alomone Labs (Jerusalem, Israel), and PDBu was from Sigma (St. Louis, MO).

In order to approximate the depolarizing input that place cells receive as an animal traverses the place field, symmetric ramp-shaped depolarizing current injections were applied via the recording electrode to CA1 pyramidal neurons at either the soma or the apical dendrite (150-250 µm from soma). Two second ramps (1 second up, 1 second down) and ten second ramps (5 seconds up, 5 seconds down) were delivered to simulate different running speeds. Current amplitude was adjusted to evoke peak frequencies between 10 and 25 Hz that resemble place cell firing as recorded *in vivo* (Hargreaves et al., 2007; Resnik et al., 2012; Bittner et al., 2015).

### Experimental design and statistical analyses

Experimental data were recorded and analyzed with Igor Pro software (WaveMetrics). Asymmetry of action potential firing with respect to the depolarizing ramp was quantified as a normalized spike ratio that is referred to here as the normalized adaptation ratio. While a ratio can be obtained by simply dividing the number of action potentials fired on the up-ramp by the number fired on the down-ramp (Harris et al., 2002), this measure cannot be applied in the case where a cell fires zero action potentials (APs) on the down-ramp. Here, we define the normalized adaptation ratio as:

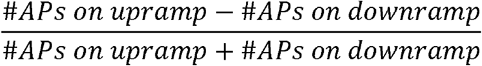

This value ranges from −1 to 1. Positive values indicate asymmetry shifted towards the up-ramp, the extreme case (a value of 1) being a cell that only fired on the up-ramp. Negative values indicate asymmetry that is shifted towards the down-ramp, the extreme case (a value of −1) being a cell that only fired on the down-ramp. A value of 0 indicates symmetric firing. Plots of instantaneous frequency versus current (f/I) use the current value at the midpoint of the interspike interval (ISI). For experiments with PDBu, changes in the rate of rise of the action potentials during a spike train were compared between conditions by comparing the time derivative of the membrane potential dV/dt (Colbert et al., 1997), to estimate the degree of attenuation of sodium conductance along the train. As most ramps evoked at least 7 action potentials, a ratio was obtained of the maximum dV/dt for the 7^th^ over the 1^st^ action potentials during the spike trains.

A typical patch clamp electrophysiology experiment with a difference in means of 1.3-2X and a standard deviation of 0.2-0.3 requires n = 6-12 for p = 0.05 and a power of 0.9 (Cohen, 1977); each experimental group in this study is made up of n ≥ 9 recording sessions. The number of cells recorded for each experiment is indicated in the Results section. Recordings were from one cell per slice; in general, a maximum of three recordings were obtained for each animal.

Statistical analyses were performed in SPSS (IBM, RRID:SCR_002865), following the tutorials and software guide from Laerd Statistics (2015, Statistical tutorials and software guides, retrieved from https://statistics.laerd.com/). Where possible, comparisons were made in the same cell before and after pharmacological treatment; the summary plots for these data include individual points and means ± standard error of the mean. For these experiments, parametric analyses (paired samples t-test and repeated-measures ANOVA) were used to compare treatments in the same neurons, since data were normally distributed as determined by the Shapiro-Wilk test for normality. Significant differences revealed by repeated measures ANOVA were followed up by post-hoc analysis using Bonferroni-corrected pair-wise comparisons. When comparing somatic and dendritic responses, data were not collected within the same neurons and displayed unequal variance. For this case, the parametric independent samples t-test, modified to accommodate unequal variance (Welch t-test) was used. The summary plot in this case consists of a box and whisker plot, showing individual data. Means ± standard error of the mean for these data are reported in the text. Differences were considered to be statistically significant when p < 0.05.

### Computational modeling

A multicompartmental CA1 pyramidal neuron model from our lab was used as a starting point (Combe et al., 2018). This model was based on the Poirazi et al. (2003a) model, with subsequent changes made in Shah et al. (2008) and Bianchi et al. (2012). The multicompartmental model from Combe et al. (2018) has 144 compartments in a reconstructed morphology (Megıias et al., 2001) (Figure 1A), where each compartment can be represented with an equivalent circuit (Figure 1B). Currents carried over from previous models include the leak conductance, an A-type K^+^ current (with different parameters for proximal and distal dendrites), a hyperpolarization-activated mixed cationic h-current, and T-type, R-type and L-type Ca^2+^ currents. Consistent with previous models, a 7-fold gradient in the h-conductance (Magee, 1998) and a 6-fold gradient in the A-type potassium conductance (Hoffman et al., 1997) were implemented along the apical trunk. In addition, in order to match the known lower amplitudes of back propagating spikes in the dendrites compared to the soma (Gasparini and Migliore, 2015), our model has a lower sodium conductance density in the dendrites compared to the soma (75% of the somatic value). The delayed rectifier in the previously cited models was replaced by separate models of K_v_1 and K_v_2 currents known to be present in CA1 pyramidal cells (Kirizs et al., 2014; Liu and Bean, 2014; Morgan et al., 2019). The K_v_1 model was taken from Zbili and Debanne (2020) and the K_v_2 model from https://senselab.med.yale.edu/ModelDB/showModel?model=184176. An inward rectifying potassium channel I_KIR_ was also added, consistent with the literature (Chen and Johnston, 2005): 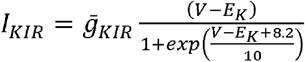. For the simulations in this work, I_M_ and I_SK_ were removed from the model for reasons explained in the text. In the previous model, intracellular calcium decayed with first order kinetics to baseline values. Calcium dynamics are not central to this study, but we incorporated the calcium balance module in Ashhad and Narayanan (2013) to make this model suitable for future extensions that will rely on calcium dynamics.

**Figure 1.**
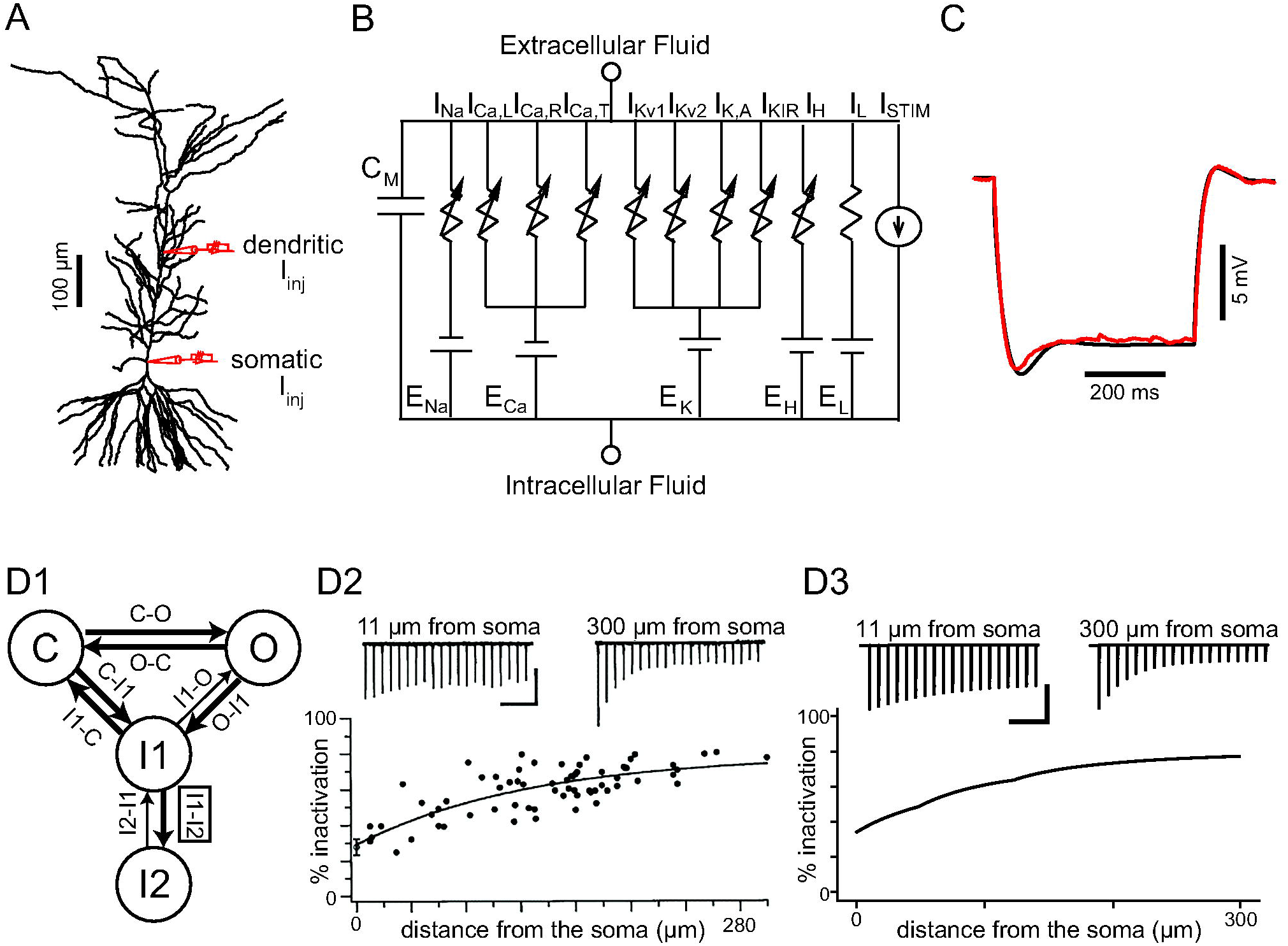
Model Schematic and Calibration. A. Morphology as implemented in the simulation package NEURON, showing the location of simulated current injection (red electrodes). Stimulus current was injected either in the soma or apical trunk (219 µm from the soma). B. Circuit diagram for each compartment. C. Superimposed voltage traces resulting from a 500 ms, 200 pA hyperpolarizing step recorded experimentally (red) or upon simulation (black) to determine the neuron input resistance. D. Sodium current model and calibration. D1. Four-state Markov model for Na_V_1.6; transitions from the I2 to the I1 state and from I1 to O are drawn with thinner arrows to indicate they are much slower than the other transitions. The I1 to I2 transition rate (box) was calibrated to reproduce data in D2. D2. Data on percent steady state sodium channel inactivation along the apical dendrite in CA1 pyramidal cells, slightly modified with permission from Figure 3B in the complete Elsevier source (Mickus et al., 1999). D3. Simulations were calibrated to fall within the envelope of the data in D2. Scale bars in D2 and D3 are 200 ms (horizontal) and 50 pA (vertical).

To better model long-term inactivation of Na_V_1.6, the dominant somato-dendritic isoform in these cells (Lorincz and Nusser, 2010), the fast Hodgkin-Huxley-type Na^+^ current was replaced in the apical dendrites and soma with a Markov model modified from Balbi et al. (2017). The original fast Hodgkin-Huxley-type Na^+^ currents were retained in the axon and basal dendrites in the absence of any data regarding long-term inactivation in those regions. This Markov model has 4 states: 1 closed state (C), a fast inactivated state (I1), a long-term inactivated state (I2) and an open state O (Figure 1D1). Current can only flow through the open state, and transitions between initial state *i* and final state *j* are governed by voltage-dependent equations in which a Boltzmann function is multiplied by a scale factor, *R_ma_*_x_, specific to each transition rate, that gives the maximum transition rate. *R_i_*_→*j*_ = *R*_max_ / (1+ exp(−(*V* −*V_H_*) / *V_S_*)). The *V_H_* for the C to O transition was made spatially dependent along the apical dendrite, varying linearly from 0 mV at 0 µm from the soma to 6 mV at ≥ 200 µm from the soma. As a result, the activation V_1/2_ for the Na_V_ channels were −13 mV proximally and −20 mV distally, consistent with data (Gasparini and Magee, 2002) showing that the V_1/2_ is more depolarized at the proximal compared to the distal dendrites.

The O-I1 and I1-O transition rates are the sum of two scaled Boltzmann functions; the parameters are reported in Table 1. To calibrate the Markov model, we simulated the experimental protocol described in Mickus et al. (1999). To replicate the cell-attached patch-clamp data therein, a simulated voltage-clamp was applied to a single-compartment iso-potential model containing only the Markov model of the Na^+^ channel. The model was held at −65 mV, and a 20 Hz train of ten 2-ms square pulses depolarized to 50 mV from rest was applied. The percentage steady-state inactivation was calculated as the difference in current amplitude between the first pulse and the tenth pulse divided by the amplitude of the first pulse. The transition rate for entry from I1 to I2 was calibrated in different spatial compartments to reproduce the observed increase in percent steady state inactivation with distance from the soma along the apical dendritic trunk as shown in Mickus et al. (1999). The following monotonically increasing expression for the maximum rate for the I1 to I2 transition produced the steady-state inactivation shown in Figure 1D3, with x representing distance from the soma in µm: for x < 49.4, *R*_max_ = 0.0299 + 0.0707(1− *e*^−^*^x^*^/126^), for 49.4 < x ≤ 124.1, *R*_max_ = 0.0091+ 0.1360(1− *e*^−*x*/126^), and for x > 124.1, −0.0933 + 0.3(1− *e*^−*x*/126^). Comparing the original data (reproduced with permission from the original Elsevier source, Mickus et al. 1999) in Figure 1D2 with Figure 1D3 shows that the percent of inactivation calculated from the simulation falls into the envelope of the experimental results.

**Table 1:**
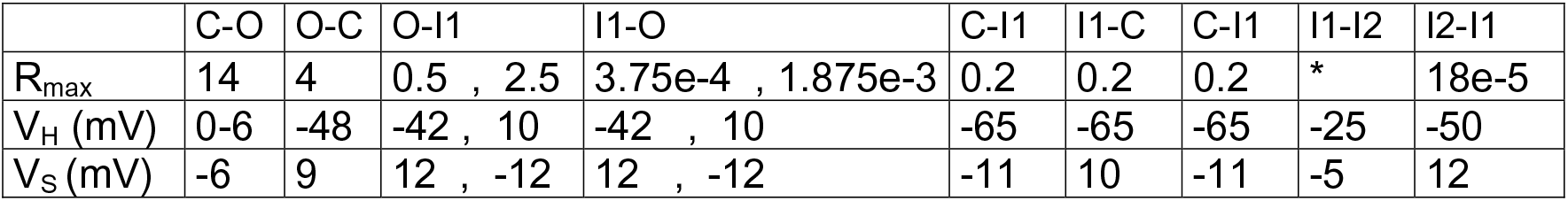
Parameters for transition rates between Markov states. The asterisk indicates a distance-dependent parameter as described in the text. *R_ma_*_x_ is in units of ms^-1^ and *V_H_* and *V_S_* are in mV.

The passive properties of the model were adjusted consistently with the literature; the response of the model neuron to a 500 ms-long square pulse of 200 pA hyperpolarizing current injection is similar to that obtained in a typical *in vitro* recording (Figure 1C). To account for the additional membrane area due to the higher density of dendritic spines at distal dendritic sites (Megı as et al., 2001), a spine factor was incorporated into the model that multiplied the capacitance and divided the membrane resistance. This factor was 1.0 in the soma, the first 40 µm of the basal tree, and the first 100 µm of the apical trunk. In the apical tree, the factor stepped to 2.0 at 100 µm, then increased with a linear dependence up to 3.5 at the point where the apical dendrite branches into the tuft (394 μ stepped to 3.5 at 40 µm from the soma. These values were based on spine densities reported by Megías et al. (2001). A sigmoidal decrease in membrane and axial resistance along the apical dendrites was implemented to further account for the decrease in input resistance along the apical dendrites (Magee et al., 1998; Poirazi et al., 2003b).

To replicate the experimental ramp protocol in the multicompartmental model, enough current was injected in the model to hold the membrane potential at −60 mV, and all transients were allowed to equilibrate before the application of the symmetric current ramp. These ramps, of two and ten second durations, as described above, were applied in the soma, and at 219 μ from the soma in the dendrites for the morphology in Figure 1A. The amplitude of the ramp was adjusted to reach similar peak frequencies as in the experiments. For the simulations, spike threshold was defined as the voltage at which the second temporal derivative of the membrane potential exceeded 20 mV/ms^2^.

### Software accessibility

Model code is freely available and can be downloaded from ModelDB at: http://modeldb.yale.edu/267140, access code: adaptationmodel

## Results

*In vivo* patch clamp recordings in rats on a linear track revealed that the synaptic input received at the soma of CA1 pyramidal cells while traversing the neuron’s place field resulted in a hill-shaped depolarization above the baseline (Epsztein et al., 2011) that generally closely followed the shape of the cell’s firing rate. Since there were no defined, repeatable shapes of this hill-shaped depolarization (Epsztein et al., 2011), in a first approximation to this spatially modulated synaptic input, we injected symmetric triangular-shaped ramps, two or ten seconds in total duration, either in the soma (Figure 2A), or in the apical trunk at ∼200 µm from the soma (between 140 and 240 µm, Figure 2B). The different durations could roughly correspond to different running speeds. For the two second ramp in the soma (Figure 2A1), there are clearly more spikes on the up-ramp compared to the down-ramp, which is quantified in the summary data of the normalized adaptation ratio (see Methods) in Figure 2C1. Larger values of normalized adaptation ratio correspond to greater asymmetry. Moreover, the inset that plots the instantaneous frequency versus the current (f/I) clearly shows hysteresis; the frequency on the down-ramp is always slower than that on the up-ramp at the same average level of depolarizing current. Concomitantly, firing initiates at a lower current amplitude than it terminates. This type of f/I plot characterizes rate adaptation (Venugopal et al., 2015). The ten second somatic ramps (Figure 2A2) show a similar pattern. The dendritic response to a two second ramp (Figure 2B1) exhibits a much more pronounced decrease in spike height during the train as well as more pronounced spike rate adaptation than the somatic ones. A Welch t-test was run to compare the normalized adaptation ratios between somatic and dendritic injections. The normalized adaptation ratio for two second dendritic ramps (0.44 ± 0.03, n = 30) was significantly higher than that for somatic ramps (0.29 ± 0.02, n = 33, t_(51.096)_ = 4.178, p < 0.001; Figure 2C1). As with the shorter ramps, there is a much more pronounced decline in spike height in the dendrites than in the soma during the ten second ramps; however, in this case a substantial recovery of spike amplitude occurs on the down-ramp (Figure 2B2). The frequency, however, does not recover on the down-ramp, indicating that these two aspects are regulated somewhat independently, as we later demonstrate. A Welch t-test was run to compare the normalized adaptation ratios between somatic and dendritic injections for ten second durations. Again, the normalized adaptation ratio was significantly higher for dendritic (0.36 ± 0.02, n = 23) than somatic ramps (0.27 ± 0.02, n = 21, t_(40.427)_ = 3.269, p = 0.002; Figure 2C2). For all groups, spiking on the down-ramp ceased at a significantly higher current value than that required to elicit spiking on the up-ramp, as illustrated in Figure 2D, which shows the difference between the current needed to initiate the last spike on the down-ramp and the current needed to initiate the first spike on the up-ramp. The mean increases, in pA, were: soma two second ramps, 22.8 ± 3.3, t_(32)_ = 6.937; dendrites two second ramps, 40.1 ± 6.6, t_(29)_ = 6.093; soma two second ramps, 22.4 ± 3.7, t_(20)_ = 6.029; dendrites two second ramps, 29.0 ± 4.4, t_(22)_ = 6.628; (p < 0.0005 for each comparison between current at last versus first spike).

**Figure 2.**
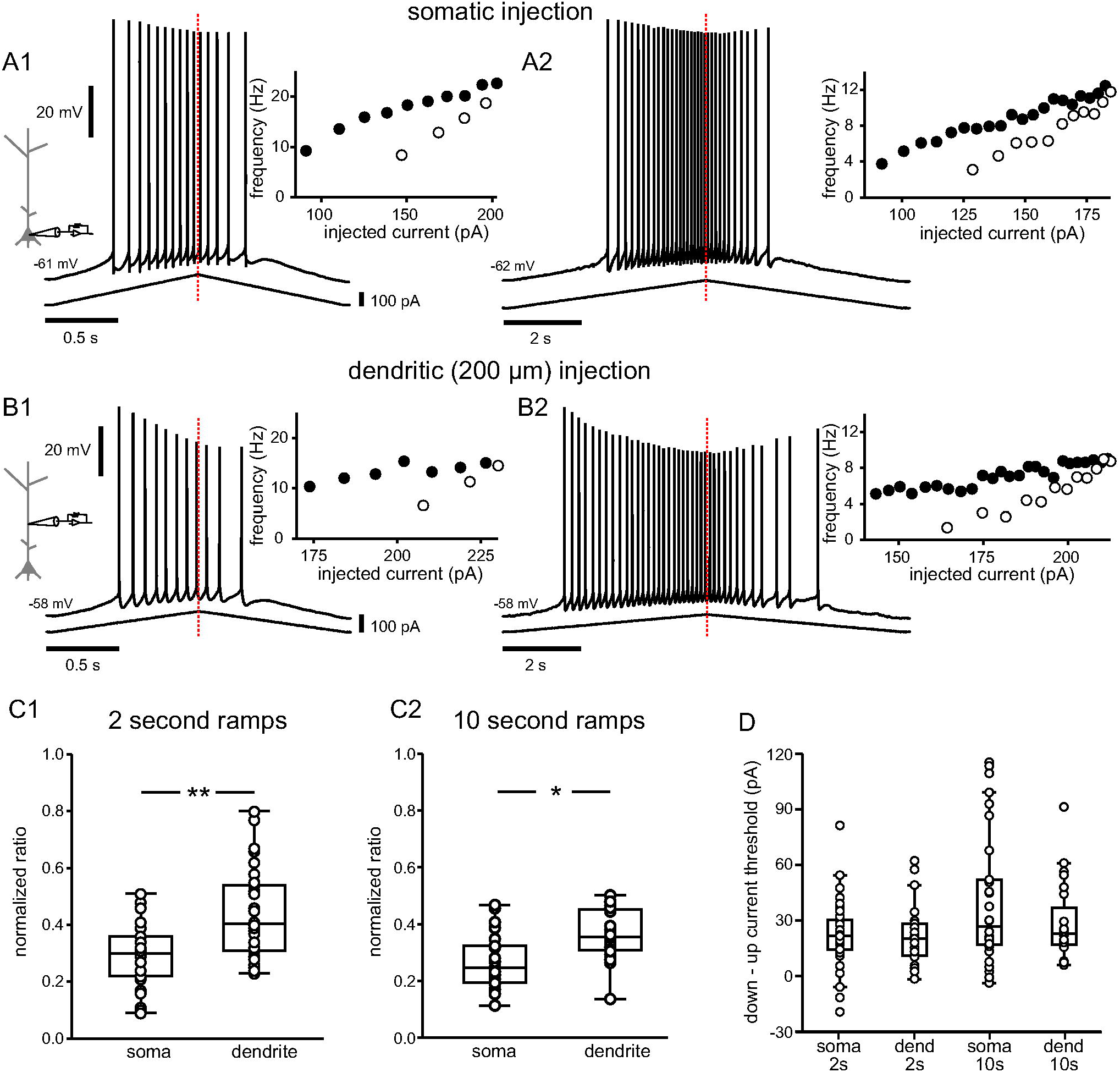
Asymmetric responses to symmetric triangular current ramps. A. Somatic injection. A1. Voltage trace recorded in the soma for a two second triangular current ramp injected in the soma. The inset shows the reciprocal of the interspike interval (ISI) plotted versus the current injected at the midpoint of the interval; filled and open circles indicate interspike intervals on the up- and down-ramps, respectively. A2. Same as A1 but for a ten second ramp. B. Dendritic injection. B1. Voltage trace recorded for a two second triangular current ramp injected in the apical dendrite at about 200 µm from the soma. B2. Same as B1 but for a ten second ramp. The red dotted lines indicate the peak of the current ramps. C1. Summary data of the normalized adaptation ratio for two second ramps for somatic (n = 33) and dendritic current injections (n = 30). C2. Summary data of the normalized adaptation ratio for ten second ramps for somatic (n = 21) and dendritic current injections (n = 23). All normalized adaptation ratios are positive, indicating more action potentials on the up-ramp. D. Difference between the current amplitude needed to initiate the first and last spike on up- and down-ramps. * p = 0.002; ** p < 0.001.

Since the SK and M-type potassium currents are likely candidates to contribute to rate adaptation, we applied pharmacological blockers of each current separately and then together, and recorded responses in the apical dendrite where adaptation is most prominent (Figure 3). We have previously shown that the SK current prevents CA1 neurons from following fast (70-100 Hz) trains of Schaffer collateral stimulation but not 40 Hz stimulation (Combe et al., 2018). In this set of experiments with current ramp injections, the SK current blocker apamin (100 nM) did not occlude firing rate adaptation (Figure 3A) and the normalized adaptation ratios were not significantly different (0.49 ± 0.05 under control conditions and 0.46 ± 0.06 in the presence of apamin, t_(10)_ = −1.511; p = 0.162, n = 11, Figure 3D). The M-type K^+^ channel blocker XE-991 (10 μM) caused the cells to fire faster at the onset of spiking (Figure 3B), but did not significantly affect the normalized adaptation ratio (0.49 ± 0.06 under control conditions and 0.52 ± 0.07 in the presence of XE-991, t_(8)_ = 0.992, p = 0.35, n = 9; Figure 3E). When both blockers were applied simultaneously, they still failed to occlude spike rate adaptation (Figure 3C), and the normalized adaptation ratio was not significantly different (0.45 ± 0.04 in control conditions and 0.50 ± 0.05 in the presence of apamin and XE-991, t_(10)_ = 1.513, p = 0.161, n = 11; Figure 3F). The voltage waveforms are different, however, in the presence of XE-991, in that the after-hyperpolarizations (AHPs) are shallower (see inset in Figure 3E and F). This may account for the initial higher firing frequency in Figure 3B and C, which is consistent with (Gu et al., 2005).

**Figure 3.**
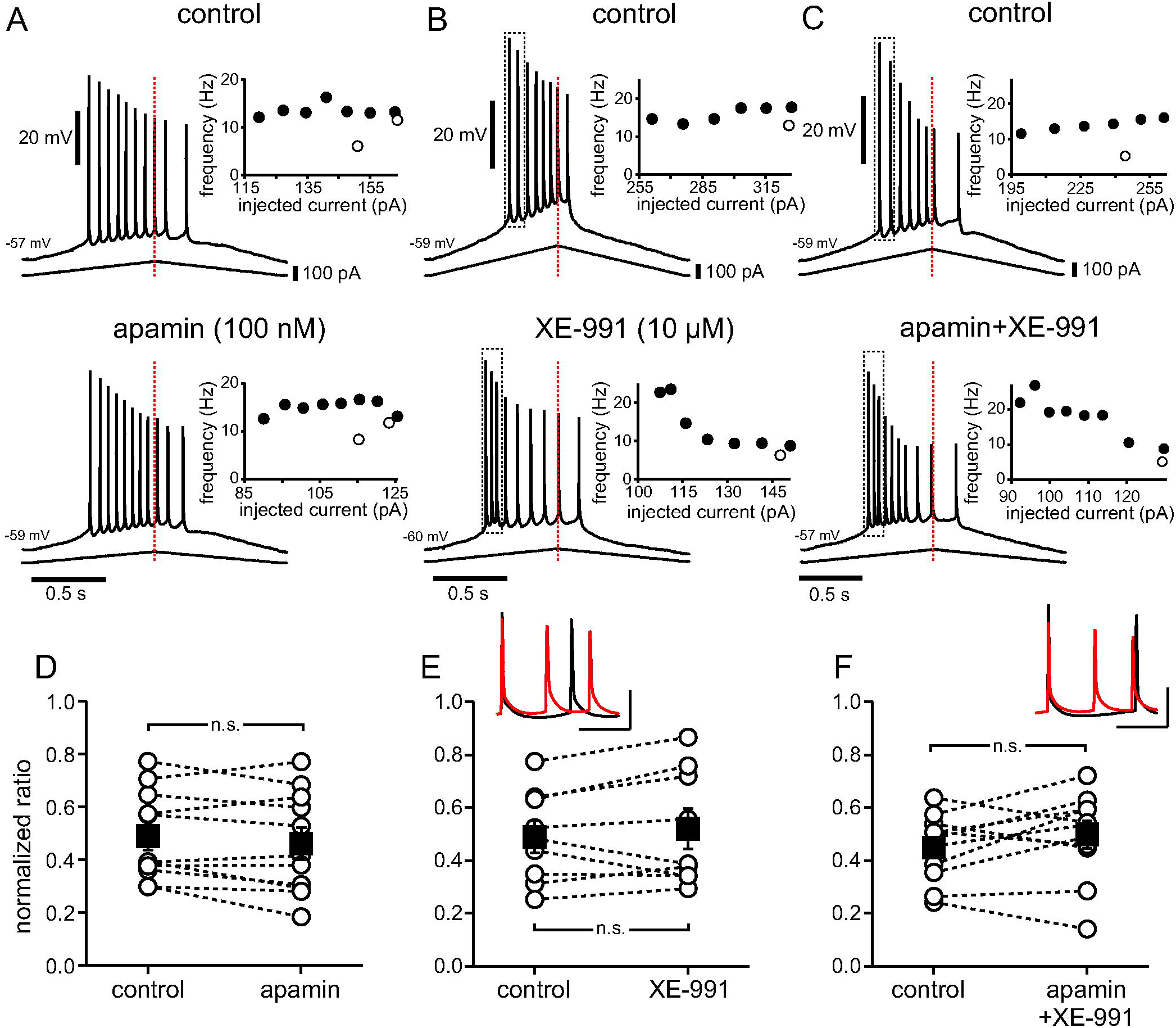
No clear effect of SK or M-type K^+^ channel block on asymmetry of ramp responses in the dendrites. Recordings were obtained at approximately 200 µm from the soma along the apical dendrite. A. Voltage trace recorded in the apical dendrite for a two second triangular current ramp (shown beneath the voltage trace) injected in the apical dendrite in control (top) and in the presence of apamin (100 nM, bottom). Insets show the reciprocal of the interspike interval plotted versus the applied current at the midpoint of the interval as in Figure 2. B. Same as A for XE-991 10 µM. C. Same as A for the combined effects of apamin 100 nM and XE-991 10 µM. D-F. Normalized adaptation ratio summary data for n = 11 neurons for apamin (D), n = 9 neurons for XE-991 (E) and n = 11 for apamin + XE (F). Dotted lines connect individual cells before and after addition of drug. Black squares with error bars represent group averages ± SEM. n.s. not significant. The insets above E and F show expanded traces (black control, red in XE-991 or apamin+XE-991), with first spikes aligned, from the areas indicated by the dotted boxes above in B and C, respectively; scalebars 20 mV vertical and 50 ms horizontal.

The observations that adaptation is more pronounced in the dendritic than in the somatic ramps, and that dendritic ramps show a prominent decrease in spike height along the train made us hypothesize that the two features could be related. The decrease in spike height during the train in the dendrites is known to be due to long-term inactivation of Na^+^ channels, also called slow, cumulative inactivation, see Colbert et al. (1997) and Jung et al. (1997). In order to add this feature to our model, we replaced the fast Hodgkin-Huxley-type Na^+^ current in the apical dendrites and soma with a Markov model modified from Balbi et al. (2017, see Methods). The previous studies in CA1 neurons often refer to slow, cumulative inactivation (Jung et al., 1997; Mickus et al., 1999). We and others (Dover et al., 2010; Navarro et al., 2020) call this process long-term inactivation instead, in order to differentiate it from a separate process called slow inactivation, which requires seconds to develop (Fleidervish and Gutnick, 1996; Ulbricht, 2005). In contrast, the long-term inactivated state is entered rapidly (within milliseconds) when fibroblast growth factor homologous factors (FHFs) bind to cytoplasmic domains of voltage-gated sodium channels (Goldfarb, 2012), but the recovery is slow, resulting in the longevity of occupancy in the long-term inactivated state. We therefore simulated the typical ramp responses recorded experimentally under control conditions in this modified multicompartmental model containing the Markov model for the Na^+^ channel; we also omitted the M-type and SK K^+^ currents, since the above results indicated these currents did not substantially promote firing rate adaptation during the triangular ramp protocol.

Figure 4A shows two and ten second simulated somatic ramps. The normalized adaptation ratio is ∼0.3 for both ramps, and rate adapting f/I curves are also observed (insets in Figures 4A1 and 4A2), thus both the voltage traces and the f/I plots are qualitatively similar to those experimentally observed in Figure 2A. The state diagram plots (Figures 4A3 and A4) show the fractional occupancy for each state of the Markov model for the Na^+^ channels, i.e. the fraction of channels that occupy a given state, as a function of time; a color diagram of the transitions is shown as an inset at the right of Figure 4A4. By definition, the sum of the fractional occupancy in all states is equal to 1. The color code at the right of Figure 4A3 orders the states in order of increasing availability to open. The least available state is the long-term inactivated (I2, magenta), since channels in this deeply inactivated state constitute a slow pool that is not readily available. Transitions from I1 to the open state O are allowed, but at an extremely low rate (see Table 1), hence the thinner arrow indicating that this rate is very small in the inset in Figure 4A4. Therefore, in general, channels must progress from the short-term inactivated state I1 (orange) through the closed state C (cyan) before opening (O, green). The occupancy in the open state during the interspike interval (ISI) is too small to be observed when plotted on the same scale as the other states, but is critical as described below. In the same way, the I1 to O transition rate, though small compared to the others, is responsible for the incomplete inactivation of the Na_V_ channel that allows for a persistent current, active during the interspike interval. For the two second ramp (Figure 4A3), occupancy in I2 accumulates throughout the spike train. For the ten second ramp (Figure 4A4), I2 recovers somewhat during the latter part of the spike train for reasons explained below. Occupancy in I1 shows a pronounced increase during each spike, but is diminished by the end of the spike train, because a large fraction of channels become sequestered in I2.

**Figure 4.**
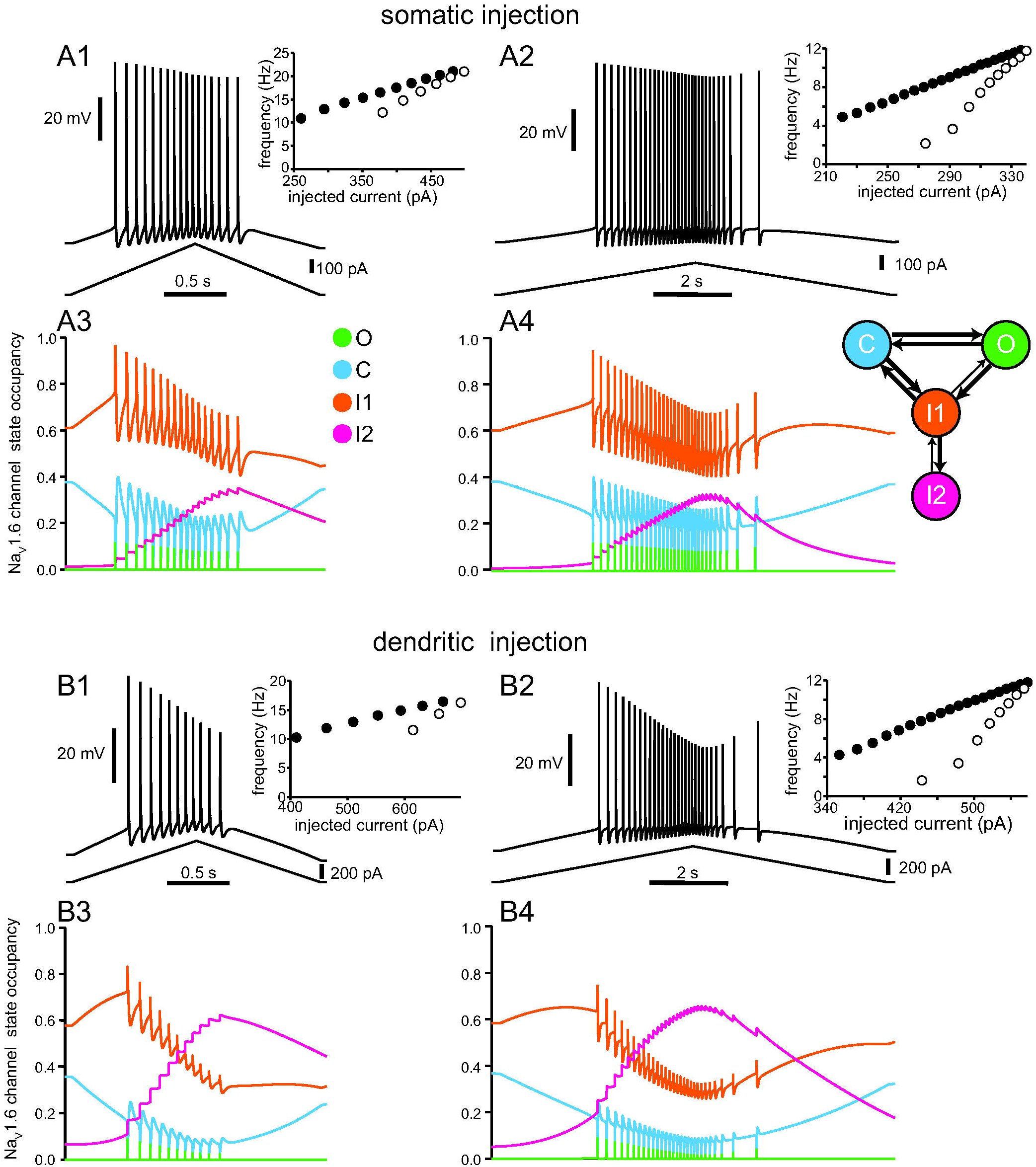
Model captures differences in somatic and dendritic responses in terms of Na_V_ availability. A. Simulated voltage traces and the resulting occupancy in each state of the Na_V_ Markov model for current ramps injected at the soma. A1. Somatic voltage response to a two second current ramp (shown beneath the voltage trace). A2. Somatic voltage response to a ten second current ramp (shown beneath the voltage trace). Insets are as in previous figures. Color plots depict the state occupancy for the corresponding two second ramp (A3) and ten second ramp (A4) and are color coded to match the Markov model schematic at right. B. Simulated voltage traces and resulting state occupancy for current ramps injected in the apical trunk at 219 µm from the soma. B1. Dendritic voltage response to a two second current ramp. B2. Dendritic voltage response to a ten second current ramp. Color plots of state occupancy (B3 and B4) correspond to the two second dendritic injection and ten second dendritic injection, respectively.

Figure 4B shows two and ten second simulated dendritic ramps. The normalized adaptation ratio is 0.45 for the shorter ramp and 0.38 for the longer ramp, and rate adapting f/I curves were again observed (insets in Figures 4B1 and 4B2). The voltage traces and f/I curves are qualitatively similar to the experimental observations in Figure 2B. For short ramps, more adaptation is observed in the dendrites than in the soma. Notably, the simulations also capture the substantially steeper decrease in spike amplitude in the dendrites compared to the soma, consistent with the experimental observations in Figure 2. In addition, for longer ramps, the simulations capture the partial recovery of spike height on the down-ramp, while the frequency keeps decreasing (compare Figure 4B2 to Figure 2B2). One important contrast between the state diagram plots for somatic and dendritic current ramps is that for somatic ramps, Na^+^ channel occupancy in I1 is always higher than in I2, whereas during dendritic ramps the I2 occupancy ends up exceeding that of I1. This is due to the spatial dependence of the parameter *R_max_* that determines the occupancy in the I2 state (see Methods and Figure 1D3), and largely accounts for the differential effects of somatic versus dendritic current injection. Thus, the more prominent effects on both spike frequency and amplitude reduction in the dendrites are due to the greater build-up of occupancy in the I2 state in the dendrites as compared to the soma (compare magenta traces in Figures 4A and B).

During the short ramps, occupancy in I2 accumulates until the end of the spike train; in contrast, during the longer ramps, occupancy in I2 peaks and then declines before spiking ceases (compare Figure 4B3 and 4B4). For these longer ramps, as the frequency slows on the down-ramp, longer interspike intervals provide more time for long-term inactivation to be removed. For sufficiently long intervals, more long-term inactivation is removed during the ISI than is added during the preceding spike, allowing occupancy in I2 to decrease. As occupancy in I2 decreases, occupancy in I1 increases, shifting a fraction of channels from the slow pool to the available pool. The spike height, which decreases during the up-ramp because long-term inactivation is accumulating, recovers to some degree on the down-ramp, as occupancy in the I2 state starts to decrease. However, the frequency does not recover; in fact, it slows down even more as the spike height recovers. Our working hypothesis was that occupancy in the I2 state is responsible for both the decrease in spike height and the decrease in frequency; therefore, it was not immediately evident how these two quantities could be differentially regulated by occupancy in the I2 state.

An explanation for these counterintuitive findings comes from the observation that, as I2 occupancy decreases and I1 occupancy increases during the down-ramp for long ramps, occupancy in C increases as well (see cyan trace in Figure 4B4). This suggests that the closed state, which constitute the readily-available pool, may play a role in the differential recovery of spike height and frequency. We looked more closely at the occupancy in various states during the longer ramps, using spike height as an additional measurement, and compared the differences between somatic and dendritic ramps by superimposing occupancy in the various states for the same levels of injected currents in the up- and down-ramps (Figure 5A-C). These plots show that occupancy in each of the states follows a history-dependent (hysteretic) trajectory. If there were no hysteresis, occupancy on the up- and down-ramps at the same level of applied current would be identical. Figure 5A shows that the occupancy in the immediately available pool, consisting of channels in the closed C state (Baranauskas and Martina, 2006), is generally lower during the down-ramp than the up-ramp at comparable values of injected current. The hysteresis is more pronounced for the dendritic injections; moreover, the occupancy in the readily available states is, in general, lower in the dendrite than at the soma (compare Figure 5A1 and A2). At spike threshold, only the Na^+^ channels in the available pool (C state) can be regeneratively recruited for the action potential upstroke. Therefore, in Figure 5D we plotted the spike height versus occupancy in the C state at spike threshold. The height was measured from threshold to peak in order to exclude contributions of the AHP to spike amplitude. Occupancy in C accurately predicted spike height, independent of whether injected current was increasing (up-ramp, filled circles) or decreasing (down-ramp, open circles), both for the soma and for the dendrites (Figure 5D). However, the dependence of the spike height on C occupancy was much steeper in the dendrites than in the soma suggesting that a smaller change in C occupancy can yield a larger difference in spike height during the spike train. Together with the larger hysteresis in occupancy states in the dendrites, this largely explains the greater decrement in spike height in the dendrites.

**Figure 5.**
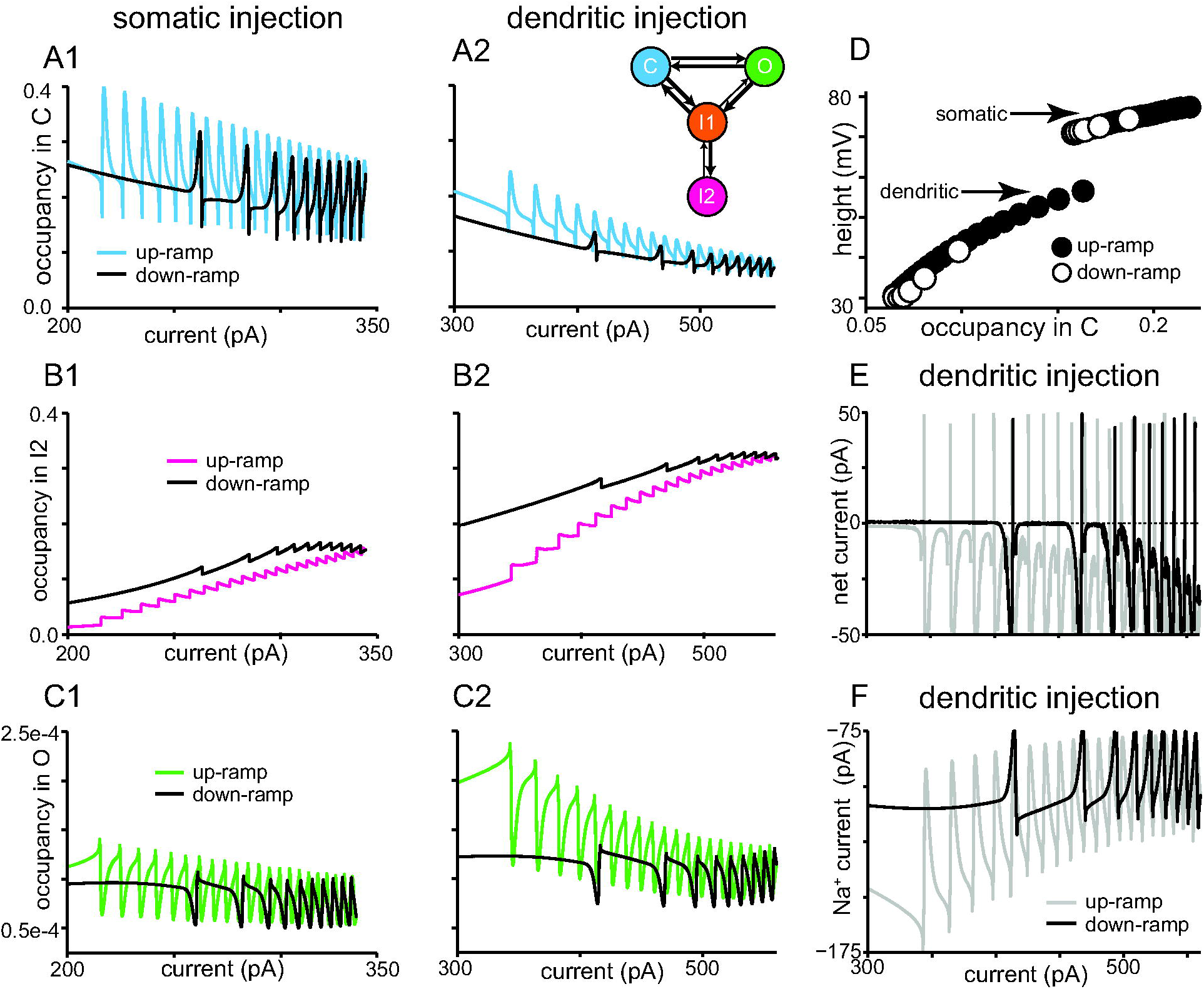
Differential regulation of spike amplitude and frequency arises from differences in Na_V_ availability during ten second current ramp. Occupancy in different Markov model states for the long ramps in Figure 4 measured during somatic injection are shown on the left and dendritic injection on the right in A, B and C. A1. Hysteresis in occupancy in the available pool of C is visible during the ten second ramp, with availability generally lower on the down-ramp (black) than on the up-ramp (cyan). A2. Hysteresis in the I2 state for dendritic injections. The inset shows the color-coded Markov model for reference. B1. Hysteresis in occupancy in the slow, unavailable pool (I2) shows that inactivation is generally higher on the down-ramp (black) than on the up-ramp (magenta). B2. Hysteresis in the I2 state for dendritic injections. C1. Hysteresis in occupancy in the open state, with higher occupancy on the up-ramp (green) than the down-ramp (black), explains the lower frequencies on the down-ramp. Values of occupancy during the spike were truncated for clarity to better show the very small open fraction during the interspike intervals. C2. Hysteresis in occupancy in the open state for dendritic injections. D. Occupancy in the available pool (C) at spike threshold is strongly predictive of spike height, measured from threshold to peak, irrespective of ramp direction (up, filled circles; down, open circles). E. Net current, obtained as the sum of the ionic currents and the injected current, as a function of injected current on the up- and down-ramp for long dendritic ramps. F. Persistent sodium current during long dendritic ramp. The persistent Na^+^ current (shown with spikes removed) is on the order of 100 pA and the decrease on the down-ramp compared to the up-ramp drives the decreases in frequency. In contrast to the occupancies, which are local values, the currents in E and F were summed over all compartments in the model.

Given the dependence of dendritic spike height on the available pool in C, it is clear that the spike height in Figure 4B2 recovers during the down-ramp because C, which is generally in equilibrium with I1, is increasing on the down-ramp (Figure 5A2, see also cyan trace in Figure 4B2). Recall that the decrease in I1 (and consequently C) is driven by the cumulative increase of occupancy in the slow pool (I2), as shown in Figure 4. We therefore looked at the occupancy in I2 for the superimposed up- and down-ramps. Again, the hysteresis is more prominent in the dendrites (Figure 5B2) than the soma (Figure 5B1). The differential regulation of the spike height and frequency by occupancy in the I2 state occurs in part because the frequency does not depend directly upon occupancy in C, which increases even as the frequency has yet to recover. Instead, the frequency depends on the very small fractional occupancy in the open state during the interspike interval (Figure 5C, with values during spikes removed). Occupancy in the open state during the interspike interval is generally lower for both soma (Figure 5C1) and dendrite (Figure 5C2) on the down-ramp compared to the up-ramp at comparable values of injected current, but again the effect is more prominent in the dendrite. Occupancy in the open state during the ISI determines the level of persistent Na^+^ current; this level, together with other factors, particularly the level of injected current, determines the frequency. The net current (Figure 5E), which is the sum of the ionic currents and the injected current and equal to the capacitive current that charges the membrane, is quite small during the ISI, on the order of < 30 pA at most, and approaches zero during flat ISIs. Although the occupancy in O is small, resulting in a small persistent Na^+^ current compared to the spiking currents, this current (Figure 5F) is large relative to the net current during the ISI. Therefore, the decrease in the Na^+^ current on the down-ramp compared to the up-ramp accounts for the slower frequencies on the down-ramp. The currents shown in Figure 5E and 5F were summed over all compartments, in contrast to the state occupancies in Figures 4 and 5A-C, which are local to the specific somatic or dendritic compartment where the current ramp was injected, as indicated. Frequency does not recover on the down-ramp because occupancy in the open state, although it increases during the down-ramp, is still below the values on the up-ramp, so the small increase is not sufficient to compensate for the constantly decreasing level of injected current on the down-ramp. A Markov model that differentiates between a fast and a slow pool as well as differentiating between a readily-available pool and a fast-inactivated pool easily accounts for this differential regulation of spike frequency and spike height, whereas we were never able to obtain these results with Hodgkin-Huxley type models. Our model describes well both spike frequency and height changes; the differential regulation of these two quantities initially appears counterintuitive and would be difficult to tease out experimentally.

In order to show unequivocally that changes in occupancy in the long-term inactivated state (I2) are sufficient to affect asymmetry and spike rate adaptation in the f/I plots, we looked at the effect of bidirectionally manipulating the levels of occupancy in I2. We initially studied the effects of removing long-term inactivation by running simulations with entry into the I2 state blocked (Figure 6A and B). These simulations confirm that firing rate adaptation in the model is indeed due to this mechanism; the f/I plots cease to show any hysteresis and become linear instead, both for somatic (Figure 6A) and dendritic current injections (Figure 6B). The model strongly suggests that spike rate adaptation is due to long-term inactivation of the sodium channels, and phosphorylation by protein kinase C (PKC) has been shown to reduce long-term inactivation of dendritic Na^+^ channels (Colbert and Johnston, 1998). Therefore, we tested the effect of reducing occupancy in the long-term inactivated state in CA1 neurons *in vitro* by applying a PKC activator, the phorbol ester phorbol-di-butyrate (PDBu, Figure 6C and D). As predicted, both somatic and dendritic ramps exhibit far less rate adaptation in the presence of PDBu compared to control. PDBu (5 µM) decreased the normalized adaptation ratio in the soma (from 0.36 ± 0.04 in control conditions to 0.13 ± 0.04, t_(10)_ = −6.738, p < 0.001, n = 11; Figure 6C and E). PDBu also prevented the reduction in spike amplitude during dendritic ramps, consistent with an effect exerted via reducing long-term inactivation of sodium channels (Colbert and Johnston, 1998). In addition, the normalized adaptation ratio for the dendritic recordings decreased from 0.50 ± 0.06 to 0.18 ± 0.05, t_(9)_ = −5.256, p < 0.001, n = 10, Figure 6D and F.

**Figure 6.**
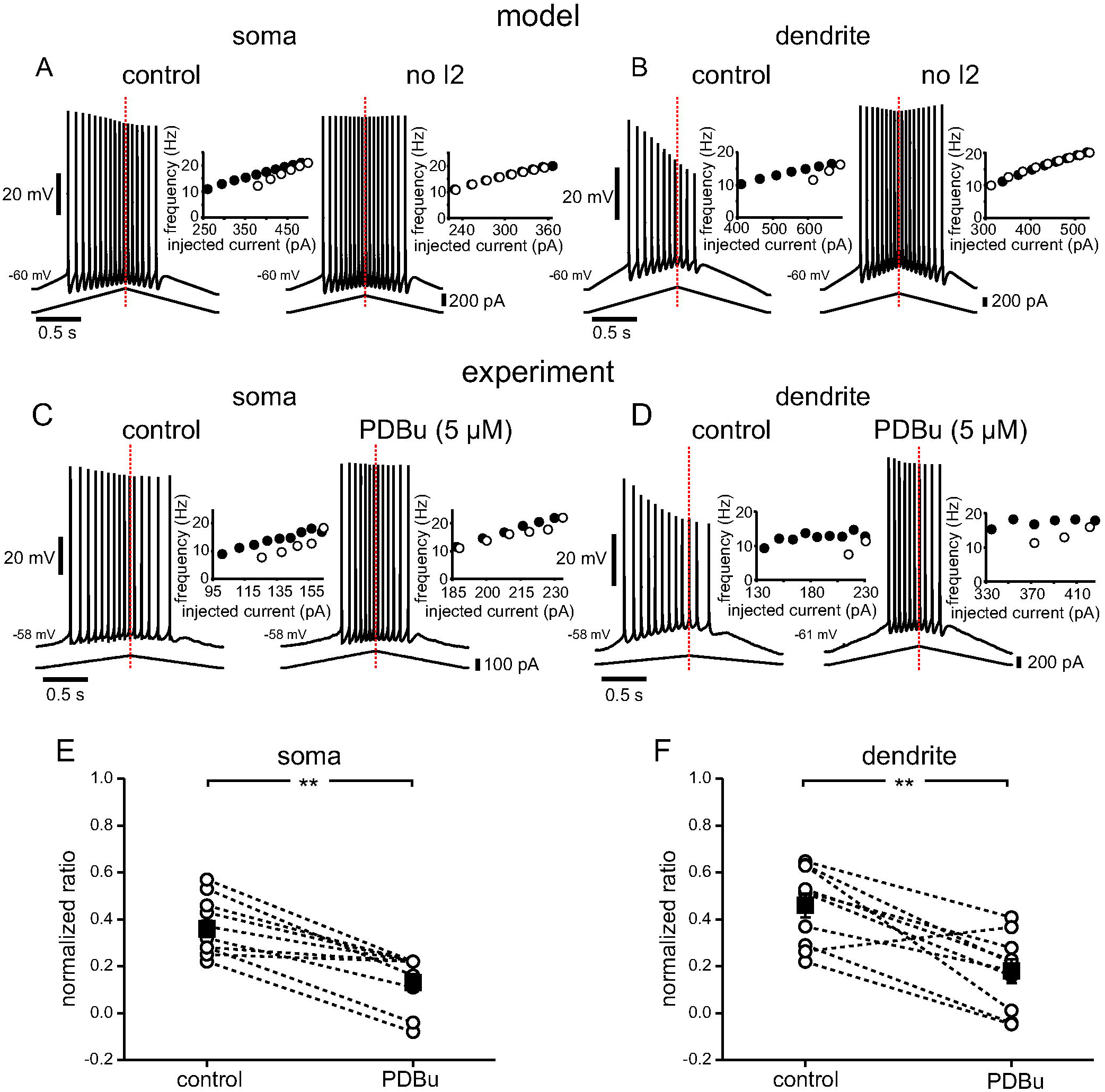
The model predicts and experiments confirm that reducing long-term inactivation reduces adaptation and asymmetry. A. Simulations of somatic current injection in control and with the I1 to I2 transition rate set to zero, meaning there is no occupancy in the long-term inactivated state. The insets show that adaptation and asymmetry are essentially eliminated by this manipulation, producing a linear f/I curve. B. Simulations of dendritic current injection in control and with the I1 to I2 transition rate set to zero. Controls in A and B are repeated from Figure 4; stimulus current was injected either in the soma or apical trunk (219 µm from the soma). C. Example experimental traces showing the voltage response to somatic current ramp injection in control and with 5 µM PDBu. D. Example experimental traces for dendritic current injection before and after addition of 5 µM PDBu. E. Summary normalized adaptation ratio data for soma (n = 11). F. Summary normalized adaptation ratio data for dendritic ramps (n = 10). Dotted lines connect individual cell averages before and after addition of drug. Black squares with error bars represent group averages ± SEM. ** p < 0.001.

Since there are other targets of phosphorylation by PKC, most notably the A-type potassium current carried by Kv4.2 channels, which is expressed at higher densities in the dendrites of CA1 neurons (Hoffman et al., 1997; Hoffman and Johnston, 1998), we performed an additional set of experiments to rule out a contribution of the A-type current to the occlusion of firing rate adaptation and asymmetry. Figure 7 shows that the specific Kv4 blocker *Androctonus mauretanicus mauretanicus toxin 3* (AmmTX3, 300 nM; Maffie et al., 2013) by itself has little effect on spike frequency adaptation or spike height (compare Figure 7B to 7A). However, the latency to fire from the ramp onset for similar current injections was found to be significantly shorter (395.0 ± 62.0 ms) in the presence of AmmTX3 than under control conditions (474.7 ± 60.7 ms; t_(8)_ = −5.788, p < 0.0005, Figure 7E). Since I_A_ is known to increase the delay to the first spike (Connor and Stevens, 1971), this results confirms a significant block of this current by AmmTX3. When added in the presence of AmmTx3, PDBu 5 µM still substantially reduced spike rate adaptation (Figure 7C), as quantified by the normalized adaptation ratio (Figure 7F), and prevented the decrease in spike amplitude during a train. The normalized adaptation ratio was significantly different across the three conditions according to a one-way repeated measures ANOVA (F_(2,16)_ = 16.100, p < 0.001). Bonferroni-corrected post-hoc pairwise comparisons found no significant changes between the normalized adaptation ratios under control conditions (0.47 ± 0.07) and in the presence of AmmTx3 (0.50 ± 0.07, p = 0.541), but a significant decrease in the presence of PDBu (0.24 ± 0.04), with p = 0.015 and 0.006, respectively, n = 9. Figure 7D shows the profile of the rate of rise (dV/dt) of the dendritic voltage traces. The maximum rate of rise corresponds to the early part of the action potential and is dependent on the available Na^+^ conductance (Fleidervish et al., 1996; Colbert and Johnston, 1998); the decrease in these peaks mirrors the decrease in spike amplitude along the action potential train evoked by the ramp. The maximum rate of rise decreased significantly along the train due to the progressively decreased availability of Na^+^ channels during the train under control conditions and in the presence of AmmTx3, but far less in the presence of PDBu (Figure 7G). A one-way repeated-measures ANOVA found that the ratio between the 7^th^ and the 1^st^ dV/dt was significantly different between the three conditions (F_(1.102,8.819)_ = 134.262, p < 0.001, n = 9). Bonferroni-corrected pairwise comparisons found no differences between control conditions (0.49 ± 0.04) and in the presence of AmmTx3 (0.49 ± 0.04, p = 1.00), but a significant increase in the presence of PDBu (0.83 ± 0.02, p < 0.001), suggesting a significant activity-dependent decrease in the fast sodium current in control and AmmTx3, that is prevented by PKC activation with PDBu. The amount of depolarization experienced during a ramp prior to the first spike does not induce substantially less inactivation in the presence of PDBu compared to in its absence. This explains the lack of effect on the rate of rise of the first (but not subsequent) spikes consistent with (Colbert and Johnston, 1998).

**Figure 7.**
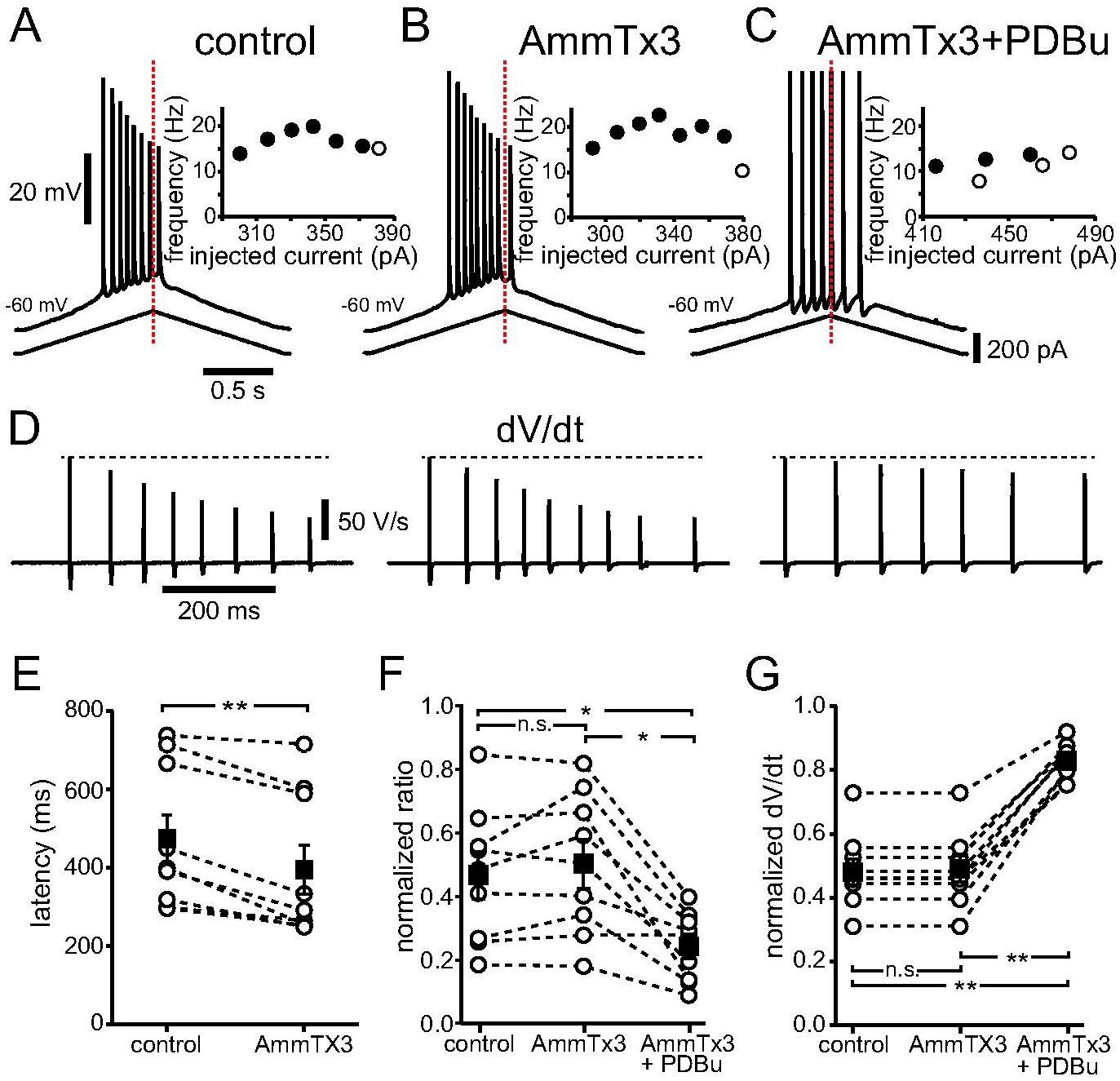
PKC phosphorylation of Kv4 channels does not mediate the reduction in adaptation and asymmetry due to phorbol ester. Top. Two second dendritic ramps. Voltage traces, current ramps and f/I curve are as in Figures 2-4. A. Control. B. AmmTx3 (300 nM) added to the bath. C. PDBu (5µM) in addition to AmmTx3. D. First temporal derivative of the voltage traces for the three conditions above. E-G. Summary data for n = 9 neurons. E. Latency to fire from onset of ramp. AmmTx3 significantly decreased the time between ramp onset and the first action potential as compared to control. F. Normalized adaptation ratios. G. Normalized peak of first temporal derivative for seventh spike. * p < 0.05; ** p < 0.0005; n.s. not significant.

In order to verify that long-term inactivation of Na^+^ channels is a generalizable mechanism to induce firing adaptation and asymmetric response to input ramps, we ran further simulations with two different morphologies (Figure 8). As shown in Figure 1D3, the values in the Markov model transitions that produce a spatially-dependent increase in the percentage of long-term inactivation similar to published data (Mickus et al., 1999) were determined for the c80761 morphology (from Ishizuka et al., 1995, Figure 8A) and the pc1a morphology (from Megías et al., 2001, Figure 8D). When this Markov model was used to the replace the original description of the Na^+^ channels in the soma and apical dendrites for these morphologies, it produced adaptive responses to symmetric current injections which were more evident in the dendrites than in the soma (compare the left panels of Figures 8B and 8E with Figures 8C and 8F, respectively). Moreover, the decrease in spike height for dendritic simulations, as previously described, was also observed. Both these features, the decrease in spike height and the adaptation were removed by blocking entry in the I2 state in the Markov model (Figure 8B, C and E, F, right panels).

**Figure 8.**
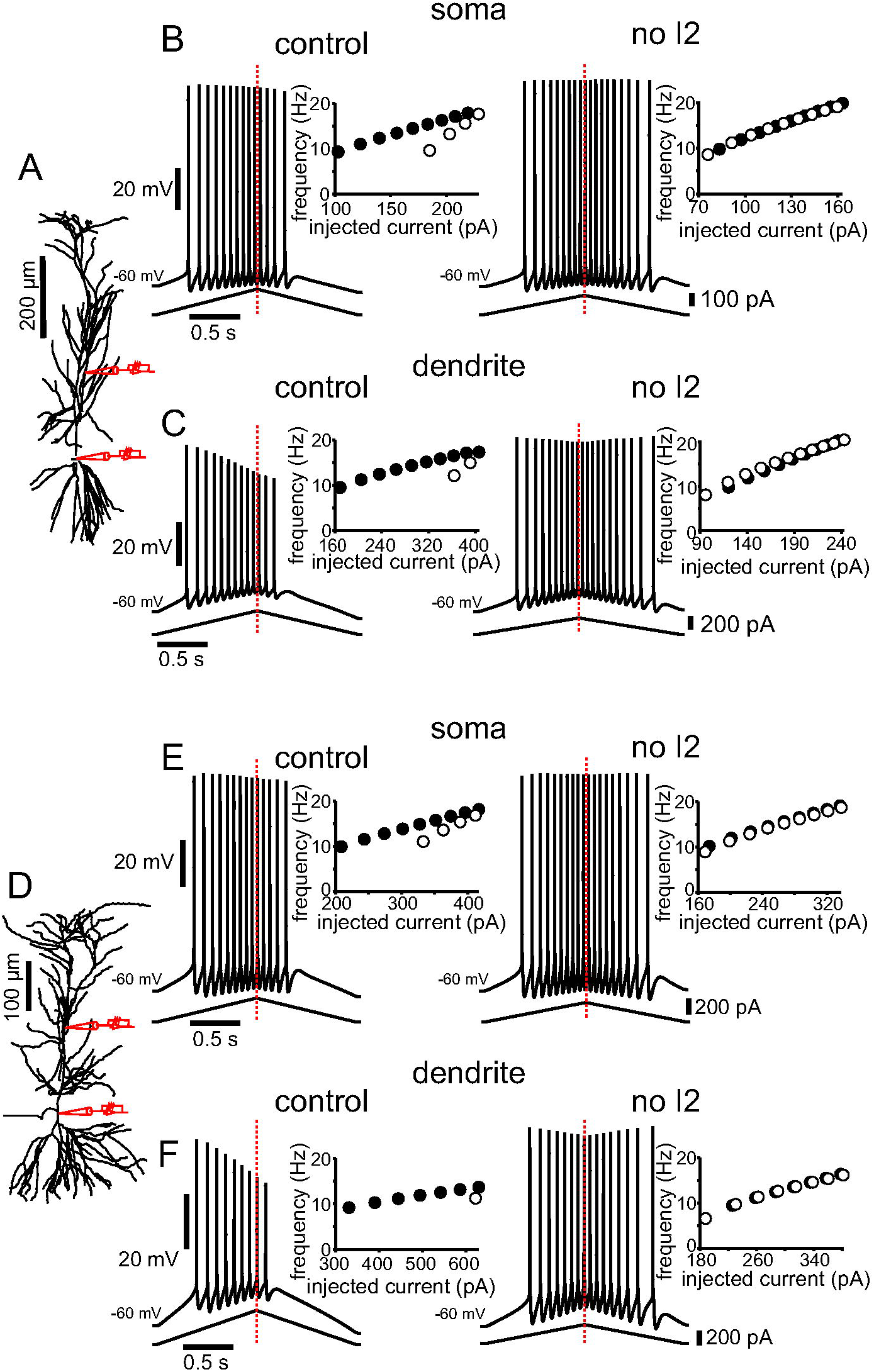
Model predictions on long-term inactivation can be generalized across different morphologies. A. Morphology (http://neuromorpho.org/neuron_info.jsp?neuron_name=c80761 from Ishizuka et al., 1995) as implemented in the simulation package NEURON, showing the location of simulated current injection. The point where the apical dendrite branches into the tuft is at 388 µm from the soma in this case. Stimulus current was injected either in the soma or apical trunk. B. Simulations of somatic current injection in control (left) and with the I1 to I2 transition rate set to zero (right). C. Simulations of dendritic current injection (at 192 µm from the soma) in control (left) and with the I1 to I2 transition rate set to zero (right). D-F same as above for morphology http://neuromorpho.org/neuron_info.jsp?neuron_name=pc1a from Megías et al., 2001. The point where the apical dendrite branches into the tuft is at 343 µm from the soma in this case. Stimulus current was injected either in the soma or apical trunk (at 208 µm from the soma). In this case, the percent of inactivation in the model (see Figure 1D3) was uniformly decreased by 5% to increase firing with dendritic injection; the resulting plot was still in the envelope of the experimental data (see Figure 1D2). The only parameters changed in the Boltzmann equations in Table 1 were the two R_max_ for the I1 to O transition (3e-4 and 1.5e-3 for A-C and 2.5e-4 and 1.25e-3 for D-F). In all cases, the insets show that adaptation and asymmetry are essentially eliminated, producing a linear f/I curve, by setting the I1 to I2 transition rate to zero.

As for the opposite scenario, in order to observe the effect of a generalized, enhanced long-term inactivation of Na^+^ channels *in vitro*, we included in the recording pipette solution a synthetic peptide corresponding to the N-terminal region (residues 2-21) of the fibroblast growth factor homologous factor 2A (FHF2A). This peptide, F2A(2-21) was shown to be necessary and sufficient to selectively induce long-term inactivation in Na_V_1.6 channels (Dover et al., 2010). When F2A(2-21) was added to the intracellular solution, the voltage traces show a prominent decrease in the spike height during the train as well an enhanced adaptation, in a manner that is clearly concentration-dependent (compare Figures 9A and B). F2A(2-21) increased the normalized adaptation ratio in a dose-dependent manner (Figure 9C; the values were 0.46 ± 0.05 for 0.25 mM, n = 8 and 0.84 ± 0.07 for 0.5 mM, n = 11). With the higher dose, frequently the neurons did not fire at all on the down-ramp, resulting in a normalized ratio of 1.0. As the drug dialyzed rather quickly into the neurons, it was not possible to obtain full control data sets; this last value of normalized adaptation ratio, however, was almost three times higher than the one recorded for somatic ramp injection under control conditions (0.29 ± 0.02), and even twice as large as the one found for dendritic ramp injections (0.44 ± 0.03, see Figure 2). To accomplish a similar situation in the model, we ran simulations where the transitions for the Markov model for all Na^+^ channels were adjusted as in the distal dendrites in Figure 1D3, therefore enhancing occupancy in I2 in the whole model neuron. When simulating injected current in the soma (Figure 9D), the resulting trace now shows a strong decrease in spike amplitude during the train, as well as a stronger adaptation, as visible from the f/I plot (the normalized adaptation ratio went from 0.31 in Figure 4A1 to 0.45 in Figure 9D). Altogether, these results support our model prediction that entry of Na_V_ channels into the long-term inactivated state promotes firing rate adaptation in response to symmetric depolarizing input.

**Figure 9.**
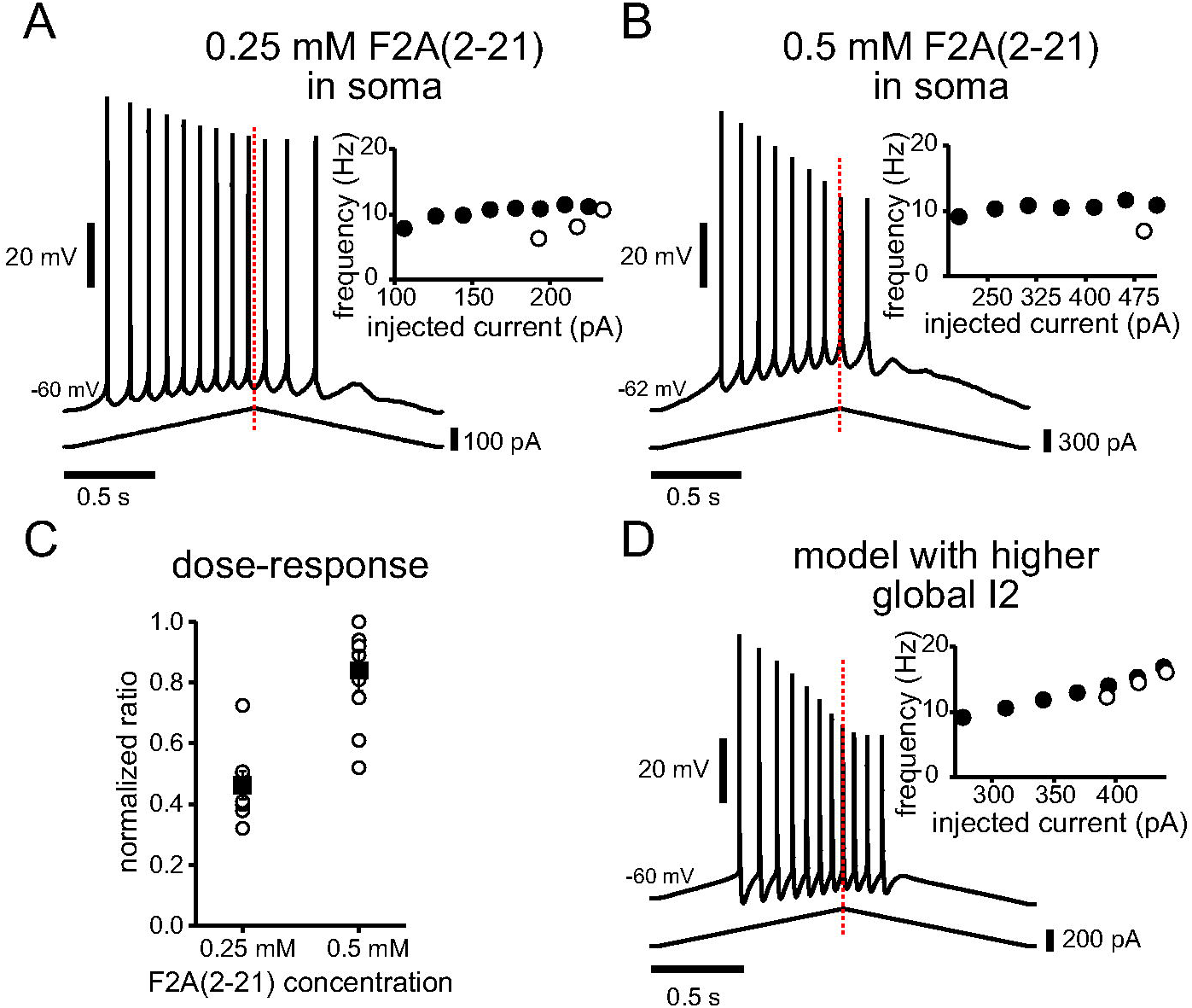
Enhancing I2 occupancy enhances adaptation and asymmetry both in the experiments and in the model. A and B. Voltage traces obtained when adding a long-term inactivation-enhancing peptide in the intracellular electrode solution (F2A(2-21)) at increasing concentrations for two second somatic current ramp injections. The peptide increases the normalized adaptation ratio and spike height decrement in a concentration-dependent manner, as shown by the average data in C (compare to the trace in Figure 2A1 with no peptide and the ratios shown for the two second somatic ramps in Figure 2C1). D. Setting I2 to a uniformly high value in the model increases the normalized adaptation ratio and results in progressively smaller amplitude spikes (compare to Figure 4A1 as a control simulation).

## Discussion

### Summary

In this study, we have shown that 1) experimentally observed adaptation in response to symmetric depolarizing ramps is more prominent in the dendrites than in the soma, 2) adaptation is not mediated by the M-type and SK K^+^ currents that mediate adaptation in other conditions, 3) adaptation is likely mediated by long-term inactivation of voltage-dependent sodium channels, and 4) a phenomenological four-state Markov model of these channels, calibrated with a distance-dependent long-term inactivation, can account for the differences in adaptation between soma and dendrites, as well as for the differential regulation of spike height and frequency by long-term inactivation.

### Significance

Since the excitatory inputs that mediate spatially-tuned firing of place cells are likely received in the dendrites, the adaptation mediated by long-term inactivation of Na^+^ channels that we observe in the dendritic ramps likely affects the position of the center of mass of place fields, with implications for spatial coding. In addition to the two and ten second ramps presented in the figures, a limited number of ramp heights and durations (one to four seconds) were also examined in this study. In general, for constant total current injection (area under the curve), adaptation is maximal for longer ramps with smaller peaks, but occurs for all shapes.

This study was performed in the context of a sustained depolarization that is thought to underlie activation of ensembles of CA1 place cells. However, the sequential activation of place cells along a linear track likely generalizes to sequential activation encoding other types of episodic memory (Wood et al., 1999), meaning that these findings may be important for understanding episodic memory in general. Also, gain control (Chance et al., 2002), defined as modulation of the slope of the frequency-current input-output relationship of a neuron, is a general feature of neural processing (Salinas and Thier, 2000). The spike frequency adaptation modulated by Na_V_ long-term inactivation is a form of divisive gain control as sustained depolarization, and the consequent adaptation, can decrease the slope of the steady-state frequency-current relationship (Fernandez and White, 2010).

### Previous work on long-term inactivation of Na_V_ channels in CA1 pyramidal cells

This work bridges studies using somatic recordings suggesting that long-term inactivation of sodium channels is responsible for frequency adaptation (Fernandez and White, 2010; Venkatesan et al., 2014) with studies showing that slow sodium channel recovery from inactivation is responsible for a decrease in amplitude during a train of dendritic back-propagating action potentials (Colbert et al., 1997; Colbert and Johnston, 1998). Our modeling work unifies these phenomena with a common mechanistic explanation of how these two processes (spike frequency and spike amplitude) are differentially regulated.

### Essential aspects of the phenomenological Markov model

Much more complicated Markov models of Na_V_ gating exist (Goldfarb, 2012; Navarro et al., 2020), but we attempted to use the simplest model possible to gain insight into long-term inactivation. We started with a published six-state model (Balbi et al., 2017) and removed one open state (Knowlton et al., 2021) and one closed state, leaving the four-state model as in Figure 1D1. Figure 10 recapitulates the main features of the model and how the various states/pools differentially regulate adaptation and spike height during the train. The first essential aspect of the model is that the I2 to I1 transition is very slow compared to that from I1 to I2, so long-term inactivation is entered rapidly but exited slowly, such that the slow pool (I2, indicated in magenta in Figure 10) sequesters a fraction of the Na_V_ channels and renders them completely unavailable for hundreds of milliseconds. Channels are recruited into this state during a spike and recover slowly during the interspike interval (see Figure 4), allowing accumulation across multiple spikes. The choice to connect I2 to the fast pool via I1 rather than O, as some models do (Goldfarb, 2012; Navarro et al., 2020), is not fundamental but phenomenological and inherited from Balbi et al. (2017). The second essential aspect of the model is that the fast pool is not monolithic but separated into three conceptual pools: the fast-inactivated (I1) pool, the readily available (C) pool and the open (O) pool. Therefore, including I2, a minimum of four pools of Na_V_ channels is required to explain our results. The closed state (indicated in cyan in Figure 10) is immediately available to be recruited into a regenerative, positive feedback loop that drives the spike upstroke, and, as shown in Figure 5D, the occupancy in this pool uniquely determines spike height. The I1 fast-inactivated state constitutes its own pool (orange in Figure 10); it is not available immediately prior to a spike but can be quickly recruited into the available pool during the after-hyperpolarization and contribute to the following spike. Finally, the open pool (indicated in green in Figure 10) has up to 20% occupancy during spiking, but in the order of 0.01% during the interspike intervals, as shown in Figure 5C. The slow pool has more occupancy on the down-ramp than on the up-ramp at comparable values of injected current, due to the history of depolarization, therefore, by a material balance, the fast pool must have less occupancy on the down-ramp. We have shown that although hysteresis in I2 drives both decreases in spike height and in frequency, these aspects are differentially regulated by the available and open pools, respectively. As shown in Figure 4, the long-term inactivated state is entered quickly during a spike and, if the subsequent interspike interval is too short to remove the amount of long-term inactivation induced by the spike, as seen during the up-ramp, inactivation accumulates. However, for sufficiently long intervals, more long-term inactivation is removed during the ISI than is added during the preceding spike, allowing occupancy in the long-term inactivated state to decrease. As a consequence, occupancy in the readily available (C) pool increases, allowing spike height to recover somewhat on the down-ramp. In contrast to the complete dependence of spike height on the available pool, frequency depends not only upon the hysteresis in the open pool, but also on the injected current. Therefore, spike height can recover to some degree on the down-ramp while the frequency is still decreasing.

**Figure 10.**
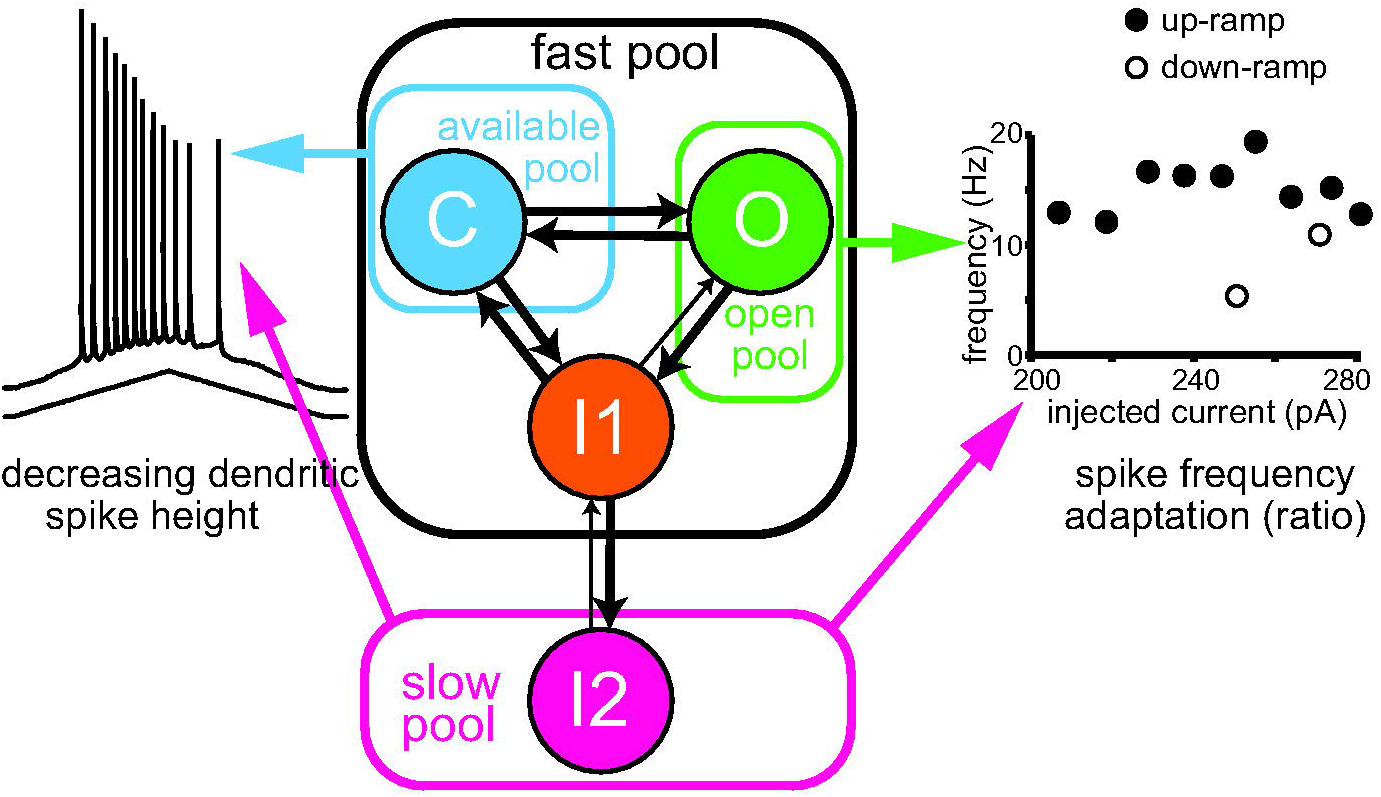
Summary diagram of the contributions of the various occupancy states to spike height and adaptation/asymmetry. The long-term inactivated state I2 causes both the decrement in spike height and spike frequency adaptation; in addition, these features are under the control of the available pool (closed state C) and the open pool (open state O), respectively. Further details are in the discussion.

### Contributions of SK and M-type potassium currents

Previous work has implicated the SK and M-type K^+^ currents in spike frequency adaptation in CA1 pyramidal cells (Madison and Nicoll, 1984; Aiken et al., 1995; Peters et al., 2005; Otto et al., 2006). Our previous work (Combe et al., 2018), showed that the SK current limited the spiking frequency of CA1 pyramidal cells in response to trains of Schaffer collateral stimulation. Action potentials failed on average for every other stimulus for > 50 Hz train in control, and blocking SK with apamin allowed the cells to follow the stimulus train more faithfully. However, at the firing frequencies (< 25 Hz) examined in this study, the SK K^+^ current did not significantly contribute to adaptation (Figure 3A). In addition, blocking the M-type K^+^ current with XE-991, allowed the initial spiking frequency to increase, but did not shift the normalized adaptation ratio in our experiments (Figure 3B and C). Similarly, linopirdine, a specific M-type channel blocker, changed the time course of spike frequency adaptation in CA1 pyramidal cells, but not the degree of adaptation attained at steady state (Fernandez and White, 2010).

### Modulation

We have shown that adaptation can be modulated by phosphorylation state (Figure 6). Since phosphorylation states depend on the neuromodulatory milieu, adaptation may be plastic, which is relevant to spatial navigation; increasing adaptation moves the center of mass of the place field in a direction opposite the direction of motion, whereas decreasing adaptation moves it in the direction of motion. In addition, since dendritic Na_V_ channels require seconds to fully recover from long-term inactivation (Colbert et al., 1997; Mickus et al., 1999), if a place field is traversed again within a few seconds of the first pass, the response will likely be attenuated. As stated above, there are also implications for gain control of the frequency-current relationship. Although we have modeled the transition rates between Markov states as parameters, in the context of neuromodulation the transition rates can be considered state variables whose values depend on the phosphorylation state of the channels.

### Generality of this mechanism of spike rate adaptation

Adaptation due to long-term inactivation of Na_V_ channels has been observed in serotonergic raphe neurons (Navarro et al., 2020). Long-term inactivation of Na_V_ channels has also recently been demonstrated to control not only adaptation, but also entry into depolarization block in different subpopulations of midbrain dopamine neurons (Knowlton et al., 2021). We conclude that long-term inactivation of Na_V_ channels may be quite generally utilized as a mechanism for spike frequency adaptation in the central nervous system.

## Acknowledgements

This work was funded by NIH R01 MH115832 under the CRCNS program to CCC and SG. We would like to thank Dr. Mitchell Goldfarb for helpful discussion.

